# WDFY4 deficiency in NOD mice abrogates autoimmune diabetes and insulitis

**DOI:** 10.1101/2022.09.02.506326

**Authors:** Stephen T. Ferris, Jing Chen, Ray A. Ohara, Feiya Ou, Renee Wu, Sunkyung Kim, Tiantian Liu, Theresa L. Murphy, Kenneth M. Murphy

## Abstract

The events that initiate autoimmune diabetes in NOD mice remain poorly understood. CD4 and CD8 T cells are both required but whether either cell initiates disease is unclear. To test whether CD4 T cell infiltration into islet required damage to β cells induced by autoreactive CD8 T cells, we selectively inactivated *Wdfy4* in NOD mice (NOD.*Wdfy4*^-/-^) using CRISPR/Cas9 targeting. Similar to C57BL/6 *Wdfy4*^-/-^ mice NOD.*Wdfy4*^-/-^ mice develop type 1 conventional dendritic cells (cDC1) that are unable to cross-present cell-associated antigens required to activate CD8 T cells. By contrast, cDC1 from heterozygous *Wdfy4*^+/-^ mice can cross-present normally. Heterozygous NOD.*Wdfy4*^+/-^ mice develop diabetes similar to NOD mice, but NOD.*Wdfy4*^-/-^ mice neither develop diabetes nor prime autoreactive CD8 T cells *in vivo*. By contrast, NOD.*Wdfy4*^-/-^ mice can process and present MHC-II-restricted autoantigens and can activate β cell specific CD4 T cells in lymph nodes, and yet do not develop CD4 T cell infiltration in islets. These results indicate that the priming of autoreactive CD8 T cells in NOD mice requires cross-presentation by cDC1. Further, autoreactive CD8 T cells are required not only to develop diabetes, but to recruit autoreactive CD4 T cells into islets of NOD mice, perhaps in response to progressive β cell damage.

## Introduction

Type 1 diabetes (T1D) is an autoimmune disease targeting insulin-producing pancreatic β-cells. Overt diabetes occurs when insulin production becomes insufficient to maintain normal glucose homeostasis. T1D in the nonobese diabetic NOD mouse strain shares many molecular, genetic and cellular features with T1D diabetes in humans (1). CD4 and CD8 T cells have both been implicated in initiating T1D. In humans, the genetic risk for developing T1D is associated predominantly with class II MHC (2), suggesting CD4 T cells may initiate disease. Class II MHC alleles, such as HLA-DQ, with mutations at position 57 of the β chain, impart the major component of susceptibility to diabetes (3) with the highest T1D association for HLA-DQ8. Similarly, in the NOD mouse, the I-Ag7 class II MHC allele is required for T1D. I-Ag7 also has a non-aspartic acid residue at position 57 of the β chain, and HLA-DQ8 and I-Ag7 exhibit similarities in their peptide binding repertoires that frequently include acidic amino acids at the peptide’s p9 residue (4;5). But CD8 T cells and MHC class I molecules are also involved. Both CD4 and CD8 T cells are required for diabetes to develop in the NOD mouse (6). MHC class I expression is required for T1D initiation and insulitis, suggesting an early role for CD8 T cells in T1D progression (7). Likewise, in humans, analysis of disease-associated alleles showed that MHC class I alleles HLA-B and HLA-A MHC contribute significantly to T1D susceptibility (8). Recent evidence in humans suggests that autoreactive CD8 T cells are present in the pancreatic T cell population in healthy individuals (9). Blocking CD8 T cell activation through T-Bet perturbation significantly inhibited T1D in several models (10). Finally, CD8 T cells are the most abundant immune cell found within diabetic human islets and many CD8 T cell MHC class I antigen epitopes have been confirmed (11;12).

Using *Batf3*^-/-^ mice backcrossed onto the NOD background, we previously reported that cDC1 are required for initiation of T1D (13). cDC1 are antigen presenting cells (APCs) that are specialized for cross-priming cytotoxic CD8 T cells to exogenously acquired antigen (14). However, we recently showed that cDC1 are also capable of priming CD4 T cells (15), so that the requirement of cDC1 for developing T1D does not indicate whether T1D in NOD mice is initiated by CD4 or CD8 T cells. We recently discovered cross-presentation by cDC1 requires the BEACH domain containing protein WDFY4 (16). Importantly, *Wdfy4* deficiency does not impair antigen processing for MHC class II presentation to CD4 T cells, providing a method to separate a requirement for general cDC1 antigen presentation from a requirement for cross-presentation to CD8 T cells. In this study, we produced *Wdfy4*^-/-^ directly in to NOD mice to evaluate the role of cross-presentation in T1D. Our results show that CD8 T cell priming by cross-presentation is required for the development of T1D in NOD mice, and that CD4 T cell priming alone is insufficient to initiate both T1D or CD4 insulitis. These results suggest that the emergence of insulitis may require progressive damage of β cells by CD8 T cells in order to recruit primed autoreactive CD4 T cells into the islet environment.

## Materials and Methods

### Mice

NOD/ShiLtJ (NOD), NOD.Cg-Tg(TcraBDC2.5,TcrbBDC2.5)1Doi/DoiJ (BDC 2.5), NOD.Cg-Tg(TcraTcrbNY8.3)1Pesa/DvsJ (8.3) mice were obtained from the Jackson Laboratory. NOD.B6-Ptprcb/6908MrkTacJ (NOD.CD45.2) mice were a gift of Dr. Emil Unanue (Washington University in St. Louis). The 8.3 and BDC 2.5 transgenic (Tg) mice were bred to NOD.CD45.2 mice to generate 8.3 CD45.2 and BDC 2.5 CD45.2 mice for T cell transfer.

### Generation NOD.*Wdfy4*^-/-^ mice

NOD.*Wdfy4*^-/-^ mice were generated essentially as previously described (16) but by directly targeting NOD zygotes in place of C57BL/6 zygotes. These sgRNAs flanking *Wdfy4* exon 4 were identified using CHOPCHOP (http://chopchop.cbu.uib.no/); *Wdfy4* gRNA1 (CATGTAGCCTTGAGGTACAT); *Wdfy4* gRNA2 (GTCCCCTTTCCTCATAGACT). Single guide RNAs (sgRNAs) were conjugated with Cas9 protein, electroporated into 0.5 day NOD zygotes and transferred into oviducts of pseudopregnant recipient mice. Offspring were screened for exon 4 deletion using PCR primers *Wdfy4* sp2 forward (GTAGGGGTCCAGTTTTGGAGG), *Wdfy4* sp2 reverse (TCCTGATCCGCGTCACTCTT) and *Wdfy4* sp1 reverse (TGGTTACACACAGCTCGTCC). One founder with complete exon 4 deletion was crossed to wild-type NOD mice and offspring intercrossed to generate experimental NOD.*Wdfy4*^-/-^ mice and controls. Mice were maintained in a specific pathogen-free facility in accordance with the Guide for the Care and Use of Laboratory Animals of the National Institutes of Health under approval by the Institutional Animal Care and Use Committee (IACUC) at Washington University School of Medicine (Assurance Number: A3381-01).

### Flow Cytometry, Antibodies and Cell Sorting

Flow cytometry was performed using a FACSCanto II or FACSAria II (BD Biosciences) essentially as described (13). Data was analyzed using FlowJo software (Tree Star Software). Pancreatic and inguinal lymph nodes (LNs) were dispersed using Cell Dissociation Solution Non-Enzymatic (Sigma-Aldrich) for 5 min at 37°C, single cell suspensions treated with 2.4G2 conditioned media (PBS, 1% bovine serum albumin, and 12.5% 2.4G2 in Iscove's Modified Dulbecco's Medium (IMDM) at 4 °C for 15 min to block Fc receptors. Antibodies included; from BD Biosciences: CD4 (RM4-5), CD8α (53-6.7), CD8β (53-5.8), CD11b (M1/70), B220 (RA3-6B2), CD19 (1D3), CD3 (145-2C11), CD45 (30-F11), Vβ4 (KT4); from Tonbo Biosciences: CD44 (IM7), CD45.1 (A20), CD45.2 (104), CD11c (N418); from Biolegend: XCR1 (ZET), Ter119 (Ter-119), Ly6G (1A8), TCRβ (H57-597), CD3 (145-2C11), CD8 (53-6.7), CD4 (RMA4-5), CD44 (IM7), CD16/32 (93), RT1B (OX-6), Vβ8.1/8.2 (KJ16-133.18); from eBiosciences: CD45.1 (A20), F4/80 (BM8). Cells were stained with fluorescent antibodies and analyzed and/or sorted via a FACSCanto II or FACSAria II (BD Biosciences). Data was analyzed using FlowJo software (Tree Star Software).

### *In vivo* T Cell Proliferation assay

The *in vivo* T cell proliferation assay was performed for BDC2.5 and 8.3 TCR Tg T cells essentially as previously described (13). Briefly, BDC2.5 and 8.3 TCR Tg mouse spleens dispersed into single-cell suspensions, washed, incubated with MagniSortTM SAV negative selection beads (Invitrogen), magnetically separated and sort-purified as B220− CD8− TCRβ+ CD4+ CD45.1+ Vβ4+ (BDC2.5) or B220− CD8+ TCRβ+ CD4− CD45.1+ Vβ8.1/8.2+ (8.3). T cells were stained with 1 μM Cell Trace Violet (CTV) (Invitrogen) for 10 min at 37 °C and quenched with 4 °C IMDM in 10% FCS, 10^6^ labeled T cells injected intravenously into recipient mice. After 3 days, draining pancreatic lymph nodes (PLNs) and inguinal lymph nodes (ILNs) were harvested, dispersed and stained with for CD45.1, CD45.2, Vβ4, 7AAD, CD4, CD44, and TCRβ (BDC2.5 transfer) or CD45.1, CD45.2, Vβ8.1/8.2, 7AAD, CD8, CD44, and TCRβ (8.3 transfer). Cells gated as CD4+ TCRβ+CD45.2+ Vβ4+CD44+ (BDC2.5) or CD8+ TCRβ+CD45.2+ Vβ8.1/8.2+CD44+ (8.3) were analyzed for CTV dilution on a FACs CANTO II.

### Diabetes Monitoring

Blood glucose was monitored daily or weekly by urine glucose readings via Diastix (Ascencia). After two consecutive readings of ≥250 mg/dL mice were considered diabetic.

### Islet Isolation and Histology

Islets were isolated as previously described (13). For histology, pancreata were isolated and placed in neutral buffered formalin for one week, paraffin embedded, sectioned, and stained with Hematoxylin and eosin (H&E).

### Statistics

Statistical analysis was performed using GraphPad Prism software version 8. Unless otherwise noted, Mann-Whitney test was used to determine significant differences between samples, and all center values correspond to the mean. P≤0.05 was considered statistically significant. Investigators were blinded to the treatments of the mice during sample preparation and data collection.

### Data Availability

The datasets generated during and/or analyzed during the current study are available from the corresponding author upon reasonable request.

## Results

### NOD.*Wdfy4*^-/-^ mice fail to cross-present β cell antigen to CD8 T cells

Deletion of the *Wdfy4* exon 4 causes splicing from exon 3 to exon 5 producing a frame shift that prematurely terminates translation (Fig. 1a-c), as previously described (16). NOD.*Wdfy4*^-/-^ mice develop cDC1 populations and other hematopoietic lineages similar to C57BL/6 *Wdfy4*^-/-^ mice (Fig. 1d) as previously described (16). We confirmed that NOD.*Wdfy4*^-/-^ cDC1 do not cross-present cell-associated *in vivo* using adoptive transfer of 8.3 TCR Tg T cells (Fig. 1e). 8.3 Tg T cells (17;18) are reactive to peptide residues 206–214 of murine islet-specific glucose-6-phosphatase catalytic subunit–related protein (IGRP) presented by H-2K^d^ (19). In heterozygous NOD.*Wdfy4*^+/-^ mice, CVT-labeled 8.3 Tg T cells proliferated in PLNs but not ILNs (Fig. 1e), confirming specific reactivity to IGRP in PLNs, but not ILNs, as expected. By contrast, in NOD.*Wdfy4*^-/-^ mice, CVT 8.3 Tg T cells failed to proliferate in either pancreatic lymph nodes (PLNs) or inguinal LNs (ILNs; Fig. 1e), indicating lack of proper cross-presentation of IGRP. These results indicate that cDC1 in NOD.*Wdfy4*^-/-^ mice cDC1 have the expected inability for cross-presentation.

**Figure 1.**
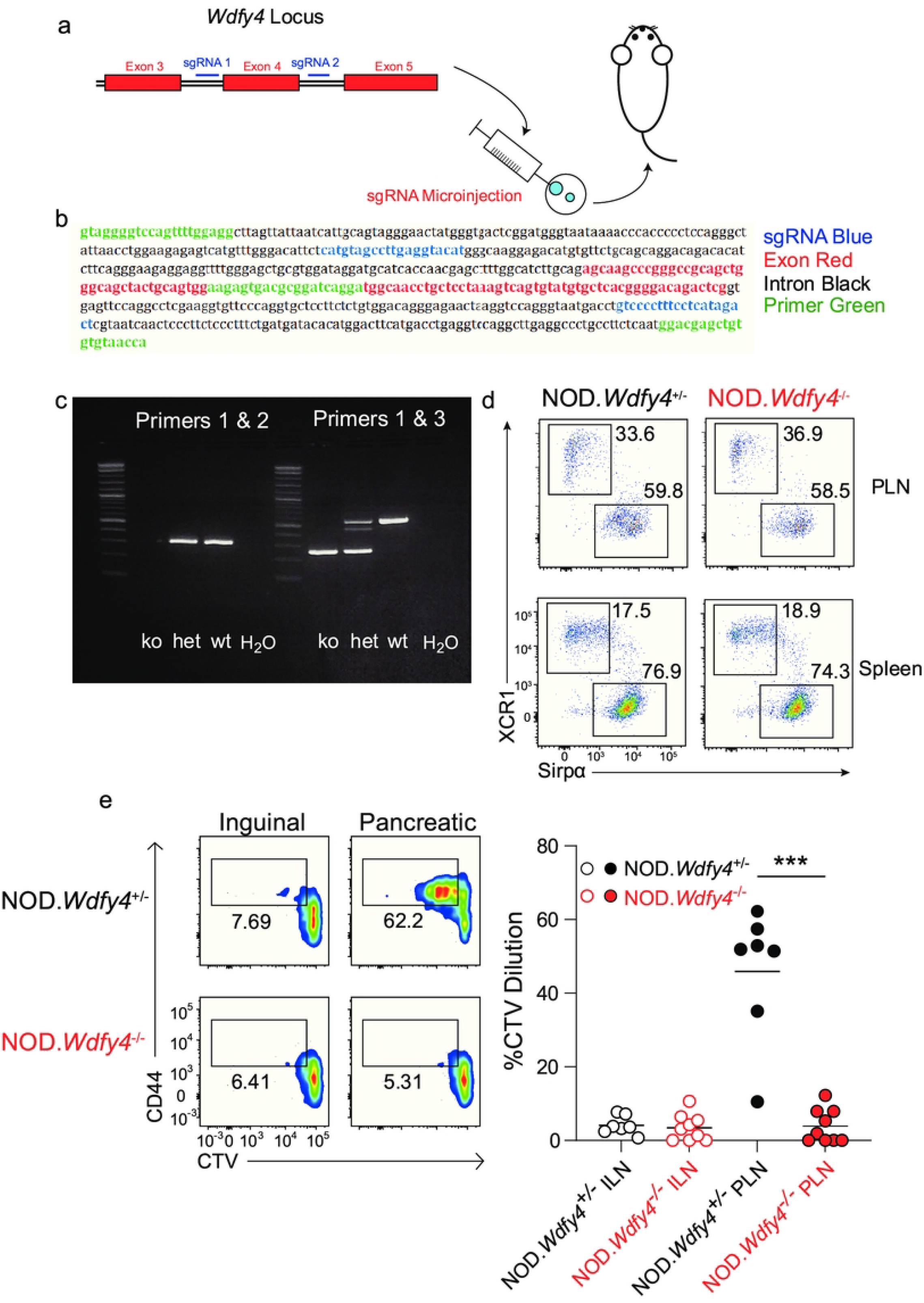
NOD.*Wdfy4*^-/-^ mice fail to prime β cell reactive 8.3 TCR tg CD8 T cells. (a) Targeting design using CRISPR Cas9 to delete *Wdfy4* exon 4. (b) Sequence showing sgRNAs, screening primers, and exons and introns for *Wdfy4* targeting design. (c) Gel of genotyping for NOD.*Wdfy4*^-/-^, NOD.*Wdfy4*^+/-^, NOD.*Wdfy4*^+/+^ mice (d) Representative flow plots of PLN (top panels) and splenic (bottom panels) cDC1 populations from NOD.*Wdfy4*^+/-^, NOD.*Wdfy4*^-/-^. Gated as B220-TCRβ+CD11c+MHCII+. (e) NOD.*Wdfy4*^+/-^, NOD.*Wdfy4*^-/-^ 6 week old female mice were injected intravenously (i.v.) with 10^6^ CTV labeled 8.3 CD45.2 cells. Left, representative flow plots of proliferating 8.3 CD45.2 T cells three days after transfer. Right, percentages of proliferating 8.3 CD45.2 cells transferred. Data are pooled biologically independent samples from three independent experiments (n=7 for NOD.*Wdfy4*^+/-^ and n=9 for NOD.*Wdfy4*^-/-^). ***P = <0.001 Mann-Whitney test.

### NOD.*Wdfy4*^-/-^ mice do not develop diabetes

We followed progression to diabetes in NOD.*Wdfy4*^-/-^, NOD.*Wdfy4*^-/+^, and NOD.*Wdfy4*^+/+^ female littermates for one year (Fig. 2). The cumulative incidence of diabetes was ~80% and ~70% in NOD.*Wdfy4*^+/+^ and heterozygous NOD.*Wdfy4*^+/-^ mice respectively (Fig. 2a). By contrast, NOD.*Wdfy4*^-/-^ females showed no progression to diabetes over the course of a year (Fig 2a). The islets of Langerhans in NOD.*Wdfy4*^-/+^ islets showed typical insulitis and peri-insulitis at both 20 weeks and 52 weeks (Fig 2b,d). By contrast, the islets of Langerhans in NOD.*Wdfy4*^-/-^ mice showed no evidence of insulitis at either 20 weeks or 52 weeks (Figures 2c, e). Thus, inactivation of *Wdfy4* completely prevents diabetes and insultis in NOD mice.

**Figure 2.**
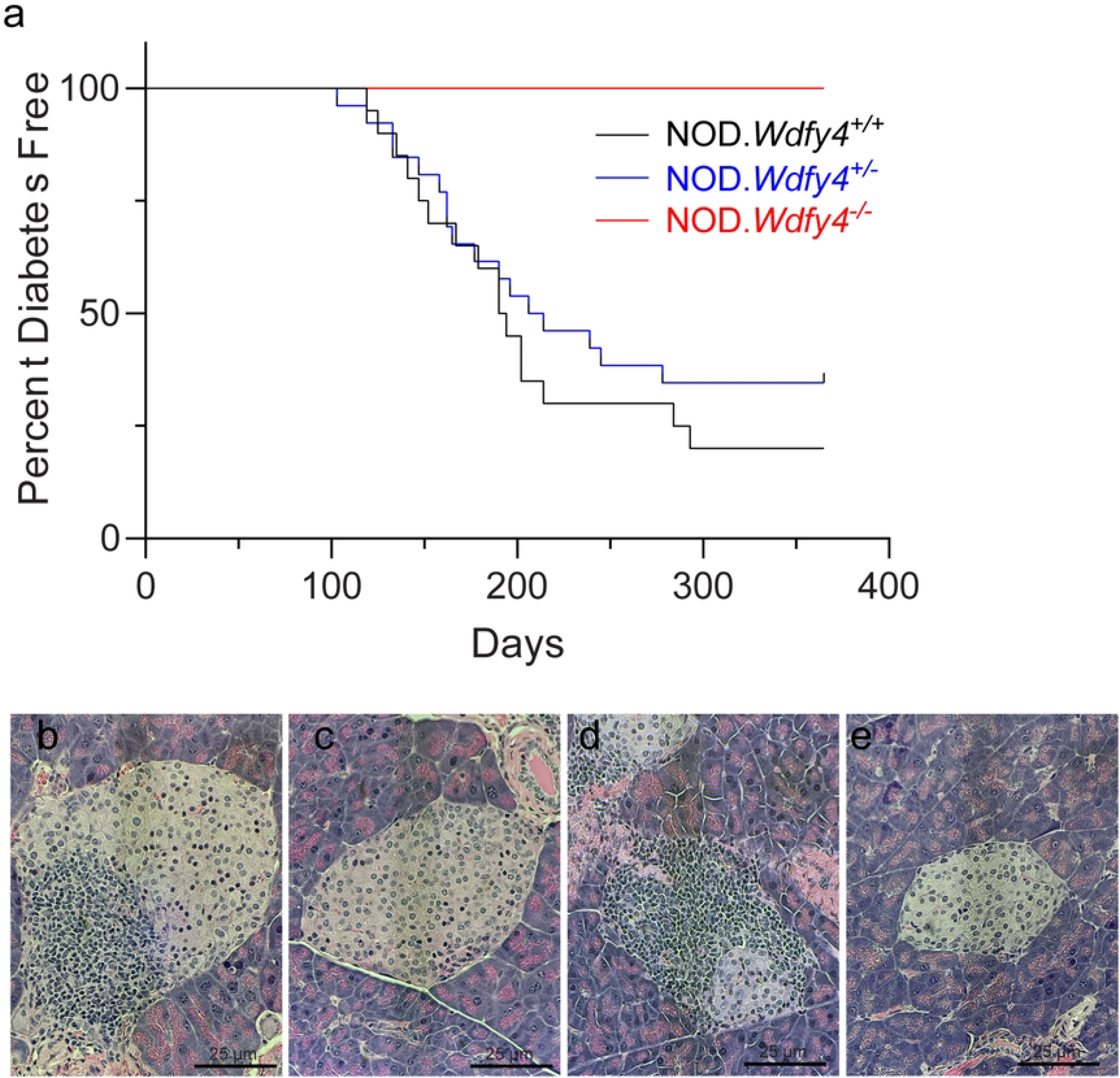
NOD.*Wdfy4*^-/-^ mice do not develop insulitis or diabetes. (a) Diabetes incidence in female NOD.*Wdfy4*^+/+^ (n=20), NOD.*Wdfy4*^+/-^ (n=27), and NOD.*Wdfy4*^-/-^ (n=17). Hematoxylin and eosin staining. (b) 20 week female NOD.*Wdfy4*^+/-^ islet. (c) 20 week female NOD.*Wdfy4*^-/-^ islet. (d) 52 week female NOD.*Wdfy4*^+/-^ islet. (e) 52 week female NOD.*Wdfy4*^-/-^ islet.

### NOD.*Wdfy4*^-/-^ mice do not develop lymphocyte infiltration into islets

We compared the immune cell infiltrates in islets of Langerhans in NOD.*Wdfy4*^+/-^ and NOD.*Wdfy4*^-/-^ mice (Fig. 3a). In 12 week NOD.*Wdfy4*^+/-^ mice, islets contained high numbers of CD45+ cells, comprising about 60% CD11c+ I-Ag7+ cells and 40% T cells (Fig 3a-c,). By contrast, in 12 week NOD.*Wdfy4*^-/-^ mice, islets contained only sparse CD45+ cells comprised primarily of CD11c+ IAg7+ cells, but very few T cells (Fig. 3 c, d). The immune compromised NOD.*Batf3*^-/-^ and NOD.*Rag*^-/-^ mice have a similar sparse CD45+ islet infiltrate comprised of CD11c+ IAg7+ islet-resident macrophages (Fig 3 a) (13). Thus, NOD mice lacking the capacity for cross-presentation lack both CD8 and CD4 T cell infiltration into islets.

**Figure 3.**
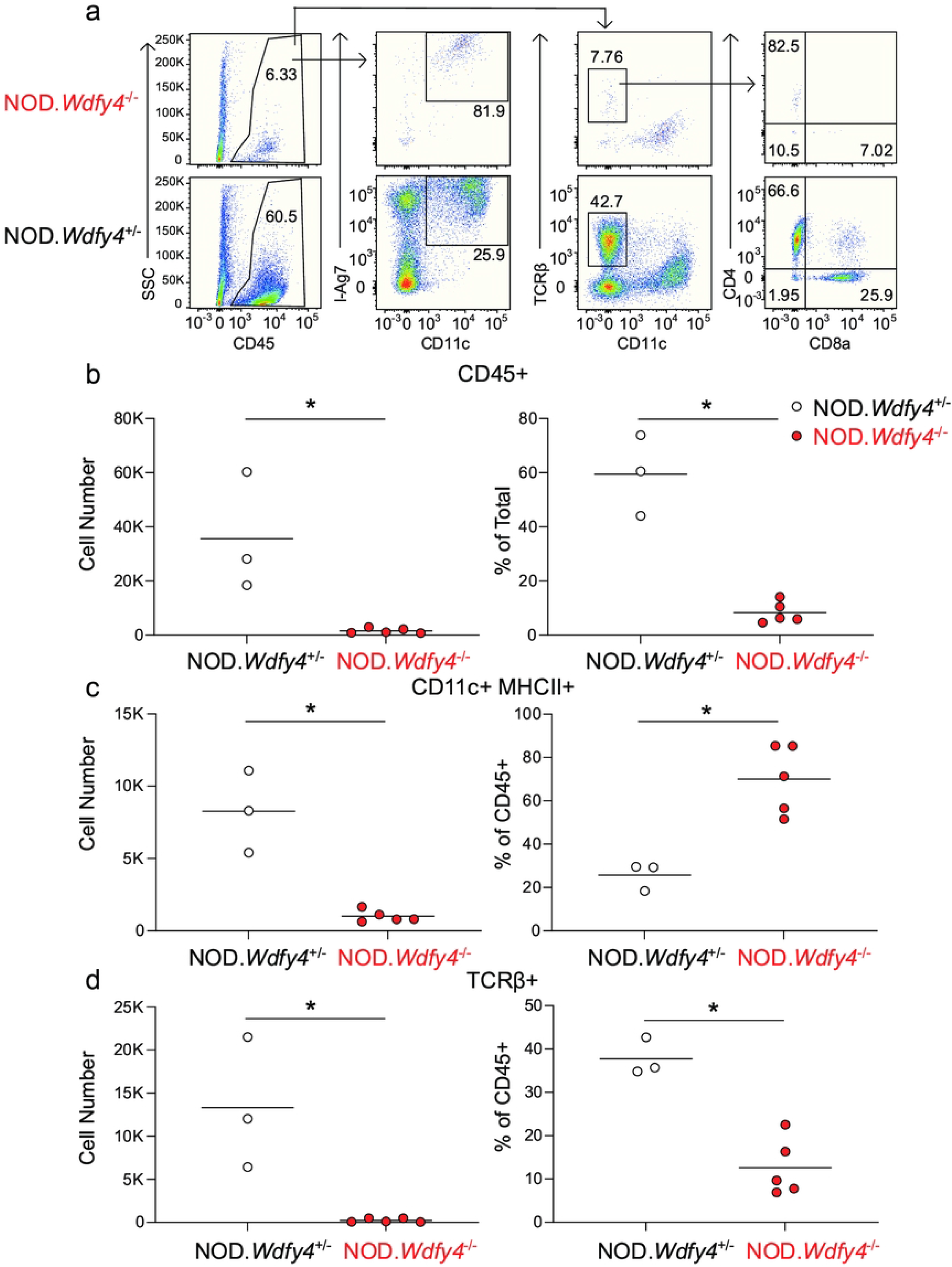
NOD.*Wdfy4*^-/-^ mice do not have inflammatory cell infiltrate. (a) Gating strategy for dispersed islets and respresentative flow plots from NOD.*Wdfy4*^-/-^ NOD.*Wdfy4*^-/-^ (top) and 12 week female NOD.*Wdfy4*^+/-^ (bottom). (b) Graph of absolute cell number (left) and percentage (right) of CD45+ cells in 12 week female NOD.*Wdfy4*^-/-^ (red) and NOD.*Wdfy4*^+/-^ (black) islets. (c) Graph of absolute cell number (left) and percentage (right) of CD11c+MHCII+ (gated as above) cells in 12 week female NOD.*Wdfy4*^-/-^ (red) and NOD.*Wdfy4*^+/-^ (black) islets. (d) Graph of absolute cell number (left) and percentage (right) of TCRβ+ (gated as above) cells in 12 week female NOD.*Wdfy4*^-/-^ (red) and NOD.*Wdfy4*^+/-^ (black) islets.

### β cell reactive CD4 T cells are activated in NOD.*Wdfy4*^-/-^ mice

BDC2.5 Tg T cells (20) recognize a β cell-specific peptide derived from chromogranin A (21). Previously, we found that BDC2.5 Tg T cells adoptively transferred into *NOD.Batf3*^-/-^ mice showed severely reduced proliferation in PLNs *in vivo* compared to WT NOD mice (20). *NOD.Batf3*^-/-^ mice lack cDC1, and so are unable to prime CD8 T cells, but may also lack an unrecognized requirement for cDC1 in MHC-II restricted antigen presentation to CD4 T cells. Alternately, the reduced BDC2.5 T cell proliferation could result indirectly from reduced amounts of antigen that may be required if the loss of CD8 T cell priming led to insufficient amounts of antigen required to drive BDC2.5 Tg T cell proliferation.

To distinguish these possibilities, we transferred CTV-labeled BDC2.5 Tg T cells to determine if autoreactive CD4 T cells could be primed by cDC1 in the absence of CD8 T cell cross-priming. We analyzed proliferation of transferred BDC2.5 Tg T cells in NOD.*Wdfy4*^+/-^ or NOD.*Wdfy4*^-/-^ mice. BDC2.5 Tg T cells transferred into NOD.*Wdfy4*^+/-^ mice proliferated in the PLN, but not in the control ILN (Fig. 4). BDC2.5 Tg T cells also proliferated after transfer into NOD.*Wdfy4*^-/-^ mice, and their proliferation was equivalent to heterozygote controls (Fig. 4). This result demonstrates that CD8 T cell cross-priming is not required for the priming of autoreactive CD4 T cells in NOD mice. Together with our results in *NOD.Batf3*^-/-^ mice, this data suggests that reduced CD4 T cell priming in *NOD.Batf3*^-/-^ mice was due to the absence of cDC1. Furthermore, we can conclude that cDC1 are the dominant cell type that primes β cell-specific BDC2.5 autoreactive CD4 T cells.

**Figure 4.**
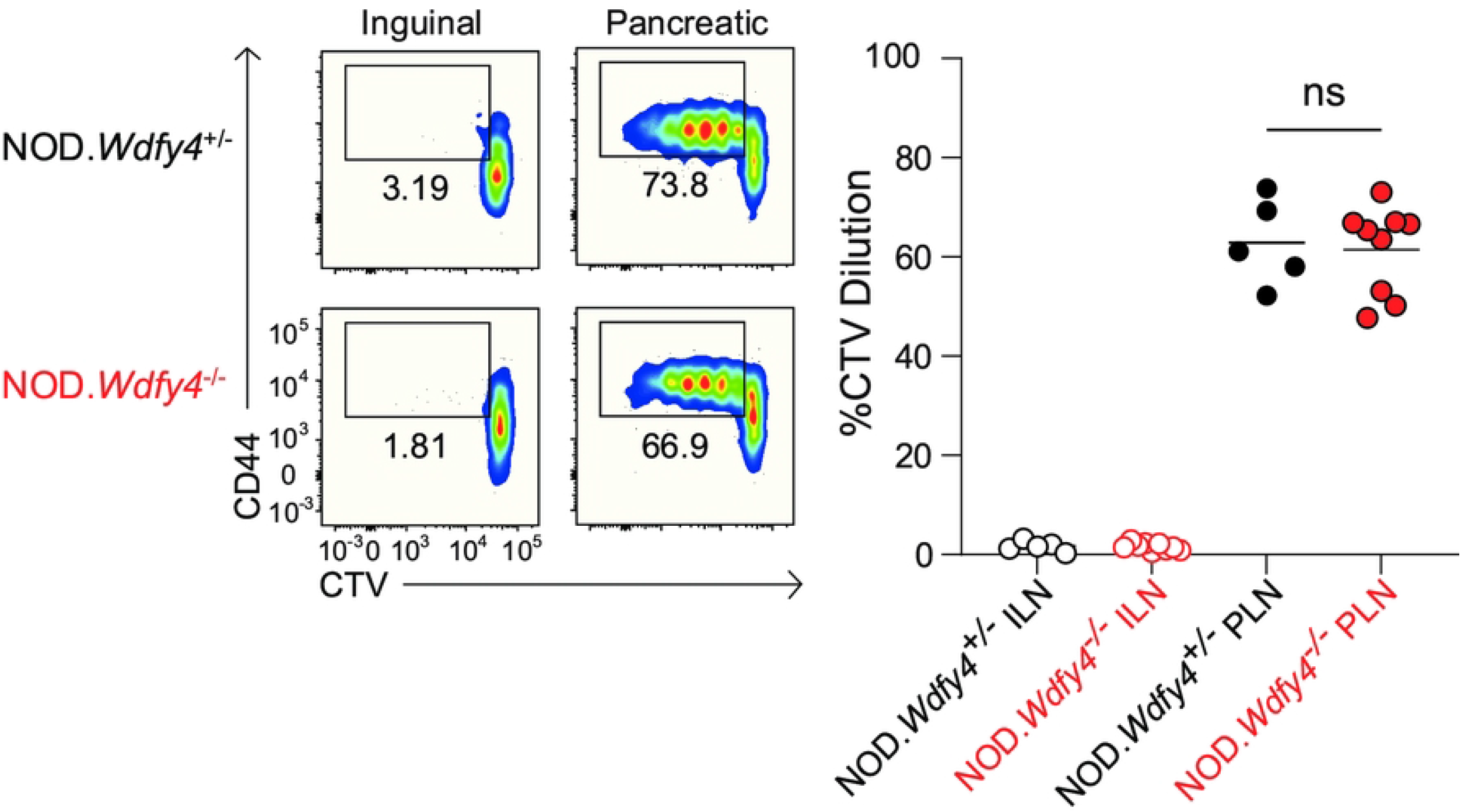
NOD.*Wdfy4*^-/-^ mice prime β cell reactive BDC2.5 TCR tg CD4 T cells. NOD.*Wdfy4*^+/-^, NOD.*Wdfy4*^-/-^ 6 week old female mice were injected intravenously (i.v.) with 10^6^ CTV labeled BDC2.5 CD45.2 cells. Left, representative flow plots of proliferating BDC2.5 CD45.2 T cells three days after transfer. Right, percentages of proliferating BDC2.5 CD45.2 cells transferred. Data are pooled biologically independent samples from three independent experiments (n=5 for NOD.*Wdfy4*^+/-^ and n=9 for NOD.*Wdfy4*^-/-^). ns = not significant Mann-Whitney test.

## Discussion

Our main finding is that selectively inactivating cross-presentation by cDC1 in NOD mice prevents activation of autoreactive CD8 T cells and averts all insulitis, but without preventing activation of autoreactive CD4 T cells. This suggests that autoreactive CD4 T cells on their own are not sufficient for causing insulitis in NOD mice. cDC1 prime CD8 T cells through cross-presentation, but are also able to prime CD4 T cells against cell-associated antigens (15). Thus, the previous finding that *NOD.Batf3*^-/-^ mice do not develop T1D could have been a result either of the loss of autoreactive CD8 T cells, or loss of autoreactive CD4 T cells, or both (13). By contrast, NOD.*Wdfy4*^-/-^ mice have a defect only in the activation of autoreactive CD8 T cells, with antigen processing for MHC-II dependent antigens and activation of autoreactive CD4 T cells being left intact. Since CD8 T cells are known to be required for T1D in NOD mice (7), the prevention of T1D in NOD.*Wdfy4*^-/-^ mice is not surprising. However, we were surprised by the complete absence of insulitis in these mice despite the maintenance of evidence for activation of autoreactive BDC2.5 Tg T cells. These results suggest the infiltration of CD4 T cells into NOD islets requires additional events beyond their initial activation in pancreatic LNs, which are likely to depend on contributions of autoreactive CD8 T cells.

Autoreactive CD8 T cells could contribute to the development of insulitis by CD4 T cells in several ways. For example, damage to β cells by cytolytic CD8 T cells may be required to recruit CD4 T cells into the islet. Indeed, recent studies using intravital microscopy have revealed that early lesions in T1D in NOD mice involve infiltration of CD8 and CD4 T cells (22), with interactions between T cells and both macrophages and DCs (13;23). Since we find that BDC2.5 Tg T cells undergo proliferation in pancreatic LNs of NOD.*Wdfy4*^-/-^ mice, it seems that cytolytic damage to islet β cells by CD8 T cells is not required for the production of islet antigens capable of trafficking to LNs. However, it is conceivable that the proliferation of BDC2.5 in pLNs observed here is not sufficient to fully induce effect CD4 T cell differentiation that is normally observed in T1D in NOD mice.

The role of cross-presentation in the development of T1D in NOD mice has been unclear. One study suggested that cDC1 are reduced in NOD mice and take on a tolerogenic activity (24). Another study suggested that the activity of cross-presentation by cDC1 in NOD mice is impaired or defective (25). In contrast, our previous results indicated that cDC1 are required for the development of T1D in NOD mice, inconsistent with a tolerogenic function (13). Secondly, our present results indicate that cross-presentation is intact in NOD mice, and in fact is required for the activation and priming of autoreactive CD8 T cells. In summary, our results suggest that full insulitis leading to T1D in NOD mice involves the coordinated activities of both CD4 T cells with CD8 T cells that are activated by cross-presentation by cDC1 in PLNs.

## Acknowledgements

This publication is solely the responsibility of the authors and does not necessarily represent the official view of the National Institutes of Health (NIH). This work was supported by the NIH (R01AI150297, R01CA248919, R01AI162643, and R21AI164142 to K.M.M., and F30CA247262 to R.W.) S.T.F. is a Cancer Research Institute Irvington Fellows supported by the Cancer Research Institute. S.T.F. is the guarantor of this work and, as such, had full access to all the data in the study and takes responsibility for the integrity of the data and the accuracy of the data analysis.

## Authors′ Contributions

**Conception and design:** S.T. Ferris, T.L. Murphy, K.M. Murphy

**Development of methodology:** S.T. Ferris, J. Chen

**Acquisition of data (provided animals, acquired and managed patients, provided facilities, etc.):** S.T. Ferris, R. A. O’Hara, F. Ou, R. Wu, S. Kim, J. Chen, T. Liu

**Analysis and interpretation of data (e.g., statistical analysis, biostatistics, computational analysis):** S.T. Ferris, J. Chen

**Writing, review, and/or revision of the manuscript:** S.T. Ferris, J. Chen, R. Wu, K.M. Murphy, T.L. Murphy

**Study supervision:** T.L. Murphy, K.M. Murphy

